# canaper: Categorical analysis of neo- and paleo-endemism in R

**DOI:** 10.1101/2022.10.06.511072

**Authors:** Joel H. Nitta, Shawn W. Laffan, Brent D. Mishler, Wataru Iwasaki

## Abstract

1. Biodiversity has typically been quantified using species richness, but this ignores evolutionary history. Due to the increasing availability of robust phylogenies, methods have been developed that incorporate phylogenetic relationships into quantification of biodiversity. CANAPE (categorical analysis of neo- and paleo-endemism) is one such method that can provide insight into the evolutionary processes generating biodiversity. The only currently available software implementing CANAPE is Biodiverse, which is written in Perl and can be used either through a graphical user interface (GUI) or user-developed scripts. However, many researchers, particularly in the fields of ecology and evolutionary biology, use the R programming language to conduct their analyses.
2. Here, we present `canaper`, a new R package that provides functions to conduct CANAPE in R. `canaper` implements methods for efficient computation, including parallelization and encoding of community data as sparse matrices. The interface is designed for maximum simplicity and reproducibility; CANAPE can be conducted with two functions, and parallel computing can be enabled with one line of code.
3. Our case study shows that `canaper` produces equivalent results to Biodiverse and can complete computations on moderately sized datasets quickly (< 10 min to reproduce a canonical study).
4. `canaper` allows researchers to conduct all analyses from data import and cleaning through CANAPE within R, thereby averting the need to manually import and export data and analysis results between programs. We anticipate `canaper` will become a part of the toolkit for analyzing biodiversity in R.

## Introduction

Quantifying biodiversity is a major goal of ecology. Quantification of biodiversity is often done on a purely taxonomic basis: counting the number of species in an area (e.g., Diamond, 1975). However, as all taxa are related to some degree by descent from a common ancestor, a thorough understanding of biodiversity is only possible by considering their evolutionary relationships. This became possible with the development of phylogenetic measures of biodiversity, such as phylogenetic diversity (PD; Faith, 1992) and phylogenetic endemism (PE; Rosauer et al., 2009) (Box 1). Such analyses are becoming much more common due to the widespread availability of robust molecular phylogenies and large spatial datasets (e.g., Mishler et al., 2020; Nitta et al., 2022; Thornhill et al., 2017).

One recently developed extension of PE is categorical analysis of neo-and paleo-endemism (CANAPE; Mishler et al., 2014). CANAPE uses phylogenetic methods to give insight into evolutionary processes that produce centers of endemism (areas with many narrow-ranged taxa; or in the phylogenetic sense, concentrations of narrow-ranged branches of the phylogeny). In theory, centers of endemism may arise via multiple processes. For example, previously widespread lineages may undergo extinction in all but a portion of their range, leading to paleo-endemism. Alternately, recently diverged lineages may only occur in a small area and lead to neo-endemism. It is also possible that a given area is home to a high concentration of both paleo- and neo-endemic lineages (mixed endemism). CANAPE involves analyzing observed patterns of PE in comparison with a null model to first infer whether an area is significantly high in PE, and then to classify those areas into centers of paleo-endemism, neo-endemism, or mixed endemism. CANAPE is widely used (> 50 publications in Google Scholar query for papers that cite Mishler et al. (2014) and mention “CANAPE”), and is a central component of the field of spatial phylogenetics.

Despite the popularity of CANAPE, it has so far only been implemented in one software package, Biodiverse (Laffan et al., 2010). Biodiverse is written in Perl and comprises an analytical engine and a graphical user interface (GUI). While Biodiverse is convenient for non-coders because of its GUI, many ecologists and evolutionary biologists use R for their analyses (Lai et al., 2019). Until now, an R user who wanted to conduct CANAPE analysis as part of a broader R workflow needed to first clean raw data, export it to Biodiverse, conduct PD and PE analyses in Biodiverse, then import the results back into R for further analysis and visualization. Also, while Biodiverse calculated the metrics needed to categorize endemism type, it did not actually perform this categorization (although this feature will be included in a future version of Biodiverse). Furthermore, the Biodiverse GUI alone does not support parallel processing, which is needed for large datasets. Parallel processing and automation of Biodiverse analyses is only currently possible using Perl scripts. A set of R scripts is available to call Biodiverse Perl scripts from R (https://github.com/NunzioKnerr/biodiverse_pipeline), but it is not an R package and does not conduct CANAPE within R.

Here, we present a new R package that implements CANAPE completely in R: `canaper`. We strove to make `canaper` simple to use and efficient. Parallel computing can be enabled with a single line of code. `canaper` has passed code review meeting the “silver” standards for statistical software at rOpenSci (Boettiger et al., 2015), and is verified against a large number of unit tests (>99% coverage). All results are reproducible by setting the random seed generator in R, in both sequential and parallel computing modes.

### Installation

The stable version of `canaper` can be installed from CRAN, and the latest development version from r-universe (https://r-universe.dev/).

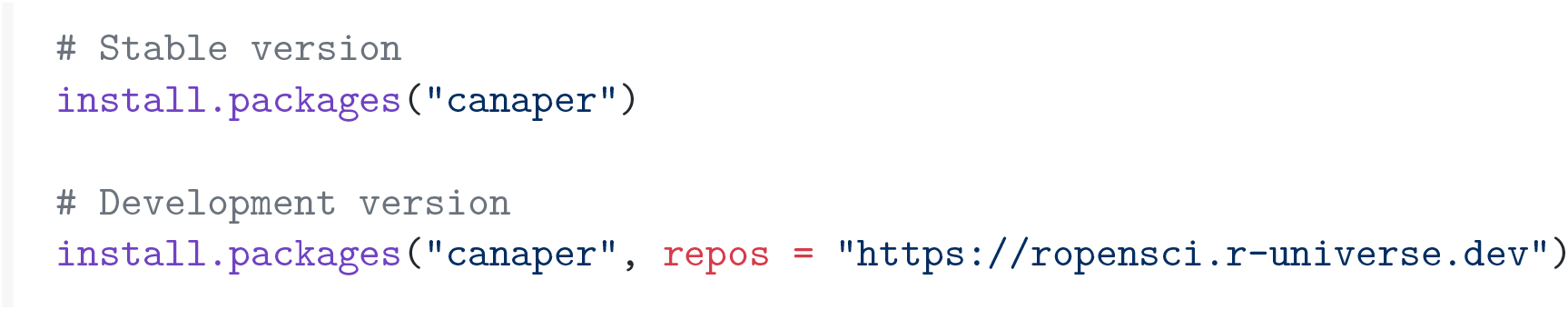

### Input data format

#### Community data

Community data is provided as a data frame or matrix, with species as columns and communities (also referred to as “sites” or “grid-cells”) as rows. In this case, the data must include both row names and column names. Community data may also be input as a tibble, in which case site names must be indicated in a dedicated column (default column name “`site`“), rather than row names since tibbles lack row names. Community data may be either presence-absence data (0s or 1s) or abundance data (integers >= 0). However, all calculations of PD and PE use only presence-absence information (i.e., no abundance weighting is used), so identical results will be obtained whether the input data is abundance or abundance that has been converted to presence-absence. Community data is typically loaded using `read.csv(), readr::read_csv()`, or other functions that can import rectangular data. The `points2comm()` function of the `phyloregion` package may be used to convert raw occurrence data (e.g., latitude and longitude of species occurrences) to a community matrix (Daru et al., 2020).

#### Phylogeny

The ape R package is used to handle phylogenies, which are stored as lists of the class `phylo`. Phylogenies should have no negative branch lengths, but are not required to be fully bifurcating. Phylogenies can be loaded with the `ape::read.tree()` function.

#### Analysis workflow

The entire CANAPE workflow can be run with two functions, `cpr_rand_test()` and `cpr_classify_endem()`. However, internally this entails several steps that the user should be aware of as follows (Fig. 1).

**Fig. 1.**
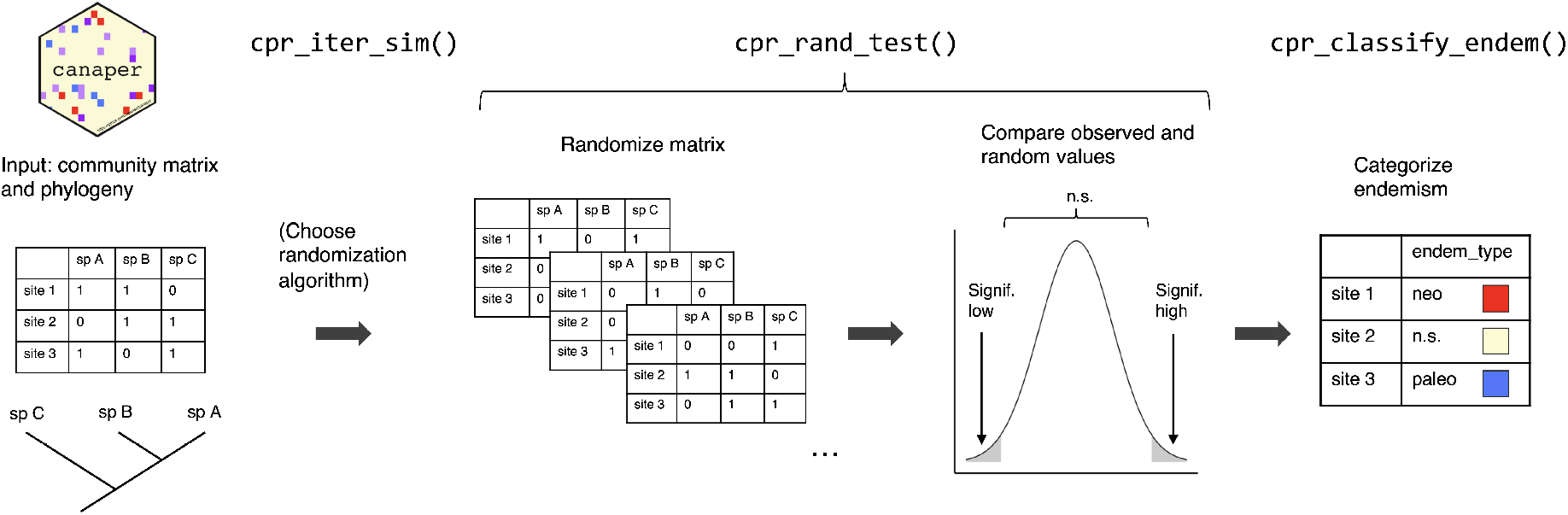
`canaper` workflow and functions to conduct categorical analysis of neo- and paleo-endemism (CANAPE). Input data are a community matrix and phylogeny. First, the user selects a randomization algorithm and its settings, with optional assistance from the `cpr_iter_sim()` function. Next, a set of random communities are generated, various metrics (e.g., phylogenetic endemism, relative phylogenetic endemism) are calculated for the input community and each random community, and the rank (*p*-value) of each observed metric is compared to those of the random communities by the `cpr_rand_test()` func-tion. Finally, endemism types are categorized on the basis of the ranked metrics with the `cpr_classify_endem()` function. For details on metrics and classification, see “Analysis Workflow”. For details on functions, see “Major Functions”.

#### Calculate observed values

First, the input phylogeny is scaled to a total length of 1 and observed phylogenetic diversity (`pd_obs`) and phylogenetic endemism (`pe_obs`) are calculated. Next, an alternative phylogeny (also referred to as the “comparison tree”) is constructed that has non-zero branch lengths set to a constant value, then rescaled to a total length of 1. PD and PE are then calculated on the alternative phylogeny (`pd_alt, pe_alt`). Relative PD and PE, the ratio of `pd_obs` to `pd_alt` and `pe_obs` to `pe_alt` respectively, are then calculated (RPD, RPE). However, the statistical significance of any of these metrics cannot be determined from observed values alone.

#### Generate random communities

PD (and by extension, PE, RPD, and RPE) will increase with taxon richness, since adding a new taxon adds new branches to the tree. To determine the statistical significance of these metrics, the observed value is compared to a distribution of values obtained from a set of random communities. The random communities are generated by a randomization algorithm that shuffles the original data.

Since the randomization algorithm influences the range of reference (expected) values, the choice of randomization algorithm is likely to have a large effect on the results. As there is no single “correct” algorithm, we have opted to provide the user with a wide range of options by implementing randomization algorithms included in the `vegan` package. `vegan`includes >30 randomization algorithms, but not all are appropriate for CANAPE. Recommended algorithms include `swap` (Gotelli & Entsminger, 2003) and `curveball` (Strona et al., 2014). These algorithms preserve the number of sites occupied by each species and the richness of each site, and both produce results comparable to one commonly used randomization algorithm in Biodiverse called `rand_structured` (see “Example: Australian *Acacia*”).

We have also provided a method for users to provide a custom, user-defined randomization algorithm using the `vegan` framework. This may be appropriate if, for example, the community matrix includes a very wide area and it is desired to restrict randomizations to subsets of the area.

#### Calculate summary statistics

Once a randomization algorithm has been selected, random communities are generated for a number of replicates set by the user, and a set of summary statistics are computed (Appendix S1). Summary statistics include the mean and standard deviation of PD, RPD, PE, and RPE of the random communities and comparisons of observed values to the random communities including standard effect size and rank, which is used to calculate *p*-values.

#### Categorize endemism

The final step in CANAPE is to categorize endemism as described in Mishler et al. (2014). Briefly, this is done by comparing significance values of summary statistics calculated in the previous step (Appendix S2). To be considered significantly endemic, a given grid-cell must first have significantly high `pe_obs` or `pe_alt` or both (one-tailed test). If this is true, the grid-cell is classified into one of three non-overlapping categories: if the grid-cell has significantly high or low RPE (two-tailed test), it is considered to be a center of paleo-endemism or neo-endemism, respectively; if RPE is not significant (but `pe_obs`, `pe_alt`, or both are), it is considered a center of mixed endemism. Centers of mixed endemism can be further divided based on *p*-value; if `pe_obs` and `pe_alt` and both significant at the *α* = 0.01 level, the grid-cell may be considered a center of super-endemism (but not all studies make this distinction).

### Major functions

#### cpr_rand_comm()

The `cpr_rand_comm()` function generates a single random community. The first argument, comm, is a community data frame (or matrix). The second, `null_model`, is the name of one of the predefined randomization to use. The remainder of the arguments are particular to specific types of randomization algorithm. `cpr_rand_comm()` is typically not called by the user directly, but is provided to help users select randomization algorithms and settings.

One feature to be aware of is that randomization algorithms in vegan are classified as either binary or quantitative. Binary algorithms are designed for binary (i.e., presence-absence) data, and quantitative algorithms are designed for quantitative (i.e., abundance) data. Either type of algorithm will accept either type of data, but binary algorithms will convert abundance data to binary and return a binary matrix (data frame).

As the calculations of PD and PE in `canaper` do not take into account abundance (i.e., no abundance weighting is used), identical results will be obtained by either using abundance data or converting abundance data to binary before analysis. In this sense, the binary randomization algorithms are appropriate for CANAPE.

The following code illustrates use of `cpr_rand_comm()` with a set of example data that comes with `canaper`, the test data from Phylocom (Webb et al., 2008).

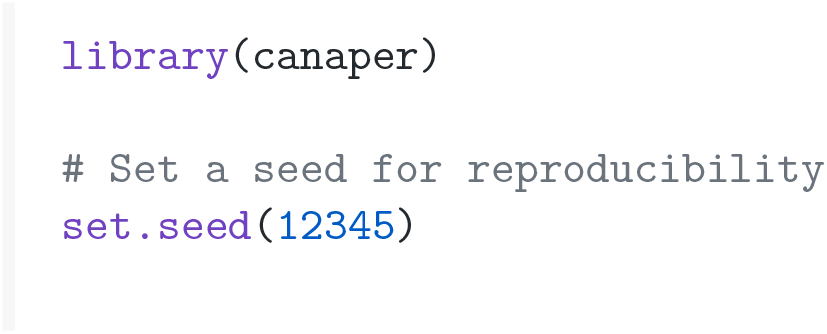
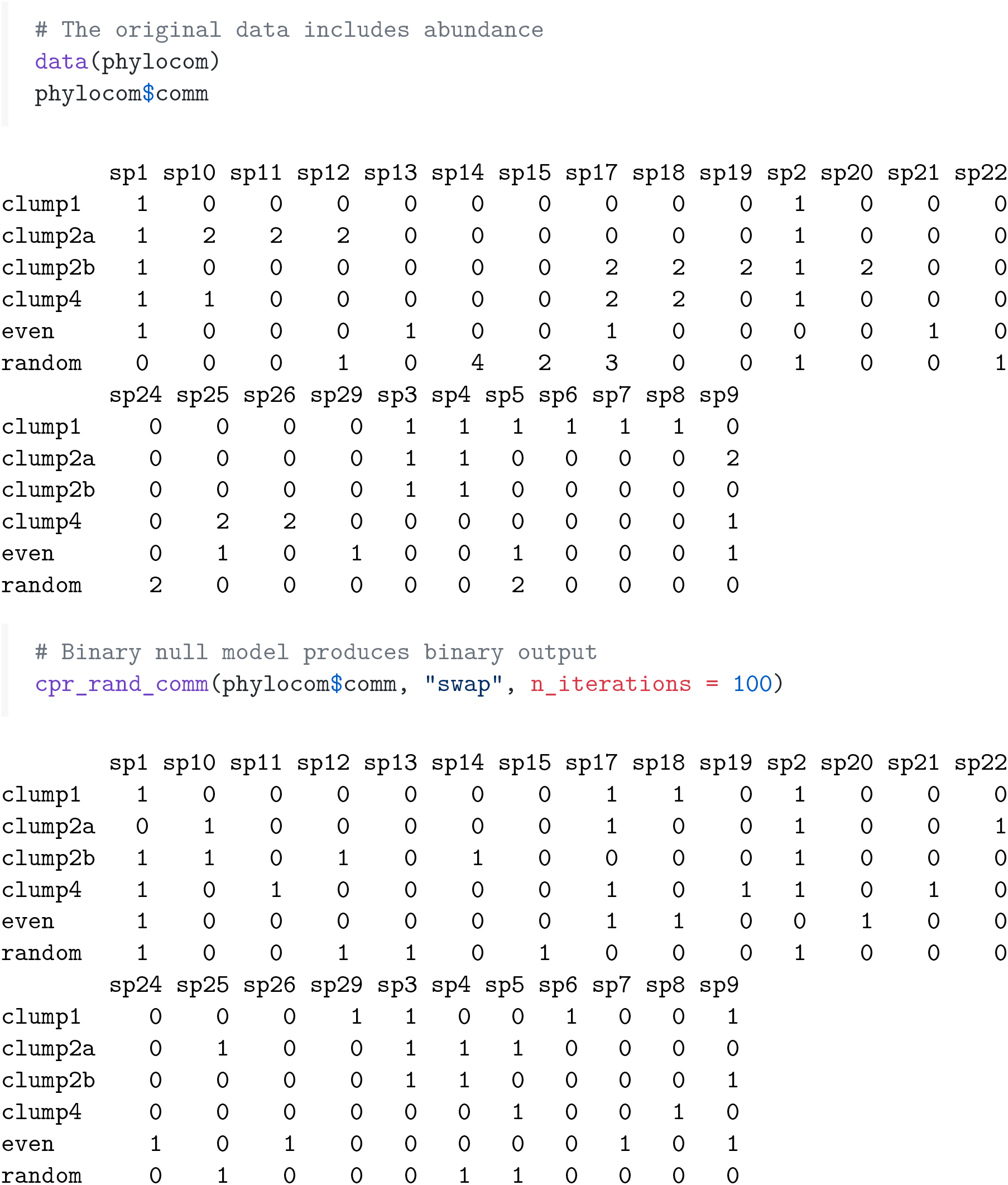

#### cpr_iter_sim()

`cpr_iter_sim()` is not required for the CANAPE workflow, but is rather a diagnostic function for use with randomization algorithms that work by exchanging cells of the community matrix such as swap (Gotelli & Entsminger, 2003) or curveball (Strona et al., 2014). Each such exchange is known as an “iteration”. It is generally impossible to know *a-priori* how many iterations are needed to completely randomize a given matrix; this number depends on properties of the matrix and the randomization algorithm. Generally, larger matrices with more skew (overabundance of zeros) will require more iterations (Miklós & Podani, 2004).

`cpr_iter_sim()` conducts successive exchanges and records the similarity between the original matrix and the randomized matrix at each iteration. The similarity values should initially decrease until an approximate minimum is reached; further iterations will only result in noise around this minimum. A number slightly larger than the smallest number of iterations needed to reach the approximate minimum value can then be used for randomizing the community. The following code demonstrates usage of `cpr_rand_test()` with the *Acacia* dataset (Mishler et al., 2014) that comes with `canaper`. The *Acacia* dataset is relatively large The Acacia dataset is relatively large (Table 1) and highly skewed (98.1% zeros).

**Table 1:** Datasets in `canaper`

| Dataset | $n$ sites | $n$ species (community matrix) | $n$ species (phylogeny) |
| --- | --- | --- | --- |
| `acacia` | 3037 | 508 | 510 |
| `biod_example` | 127 | 31 | 31 |
| `phylocom` | 6 | 25 | 32 |

The `acacia` dataset (Mishler et al., 2014) includes two outgroup taxa in the phylogeny that are not in the community matrix. The `phylocom` dataset (Webb et al., 2008) also includes more taxa in the phylogeny than the community matrix.

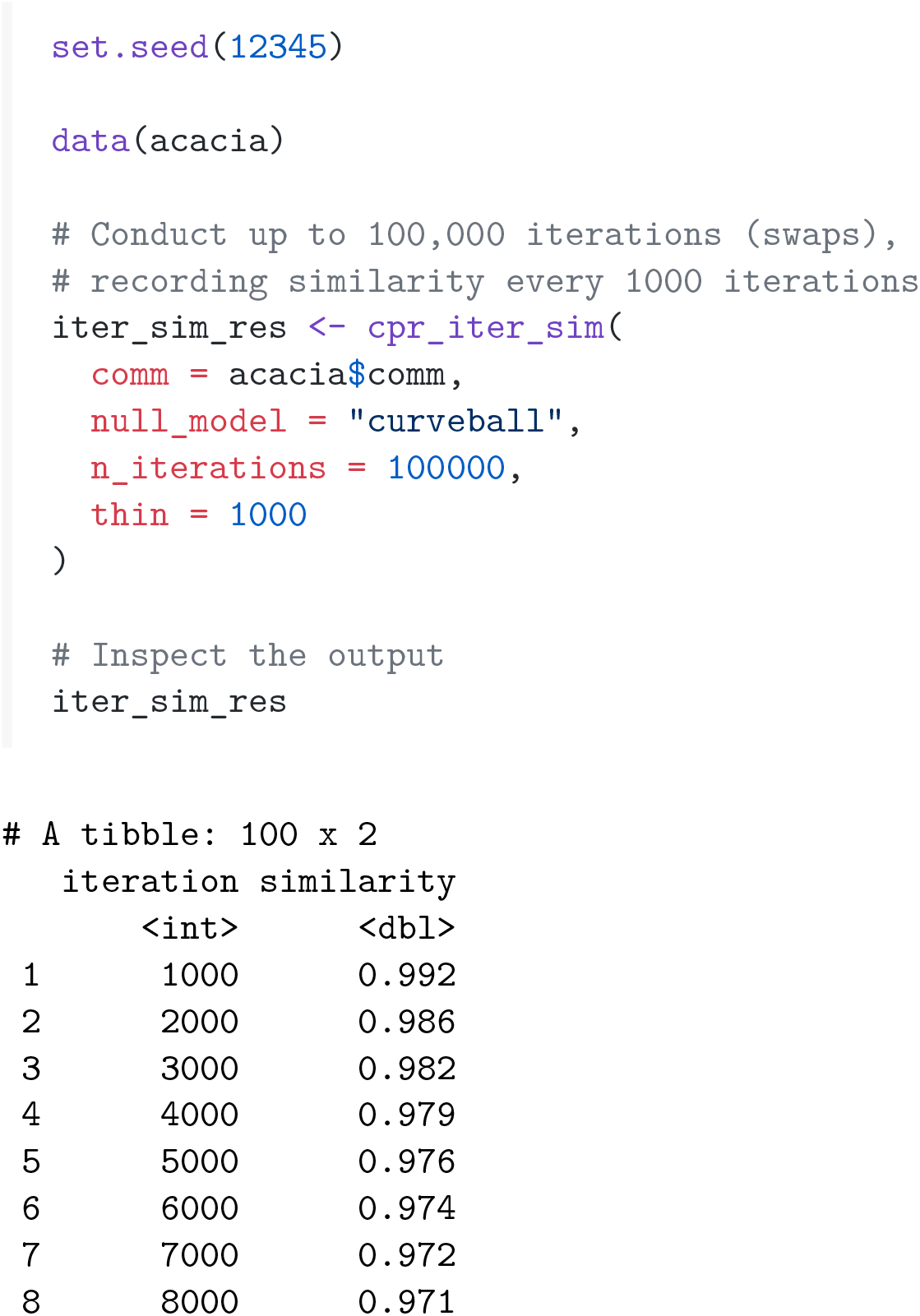
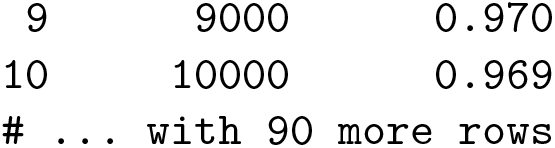

It is useful to plot the output to identify the minimum number of iterations needed. Here we do so with the `ggplot2` package.

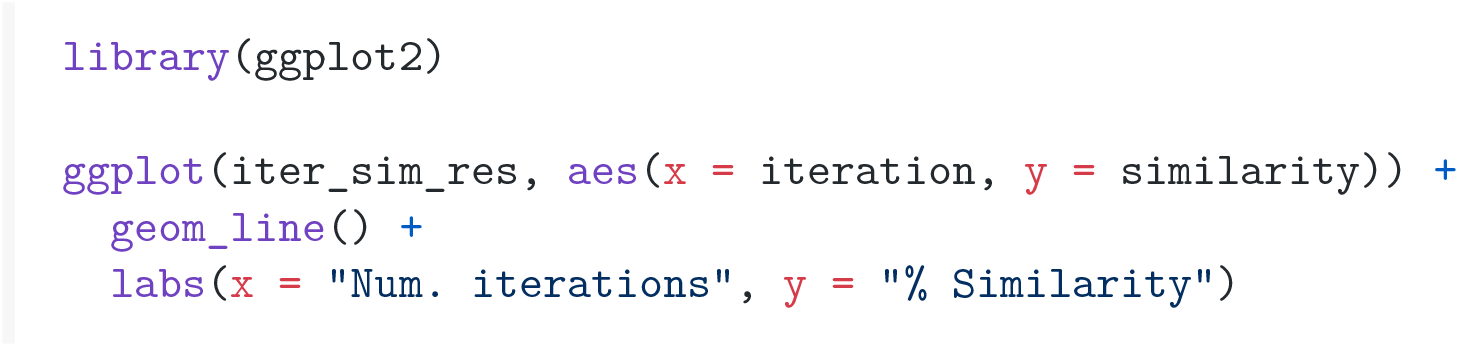

The plot shows that the community matrix becomes maximally randomized after about 40,000– 50,000 iterations (Fig. 2), and that the original matrix and randomized matrix are about 96.5% similar.

**Fig. 2.**
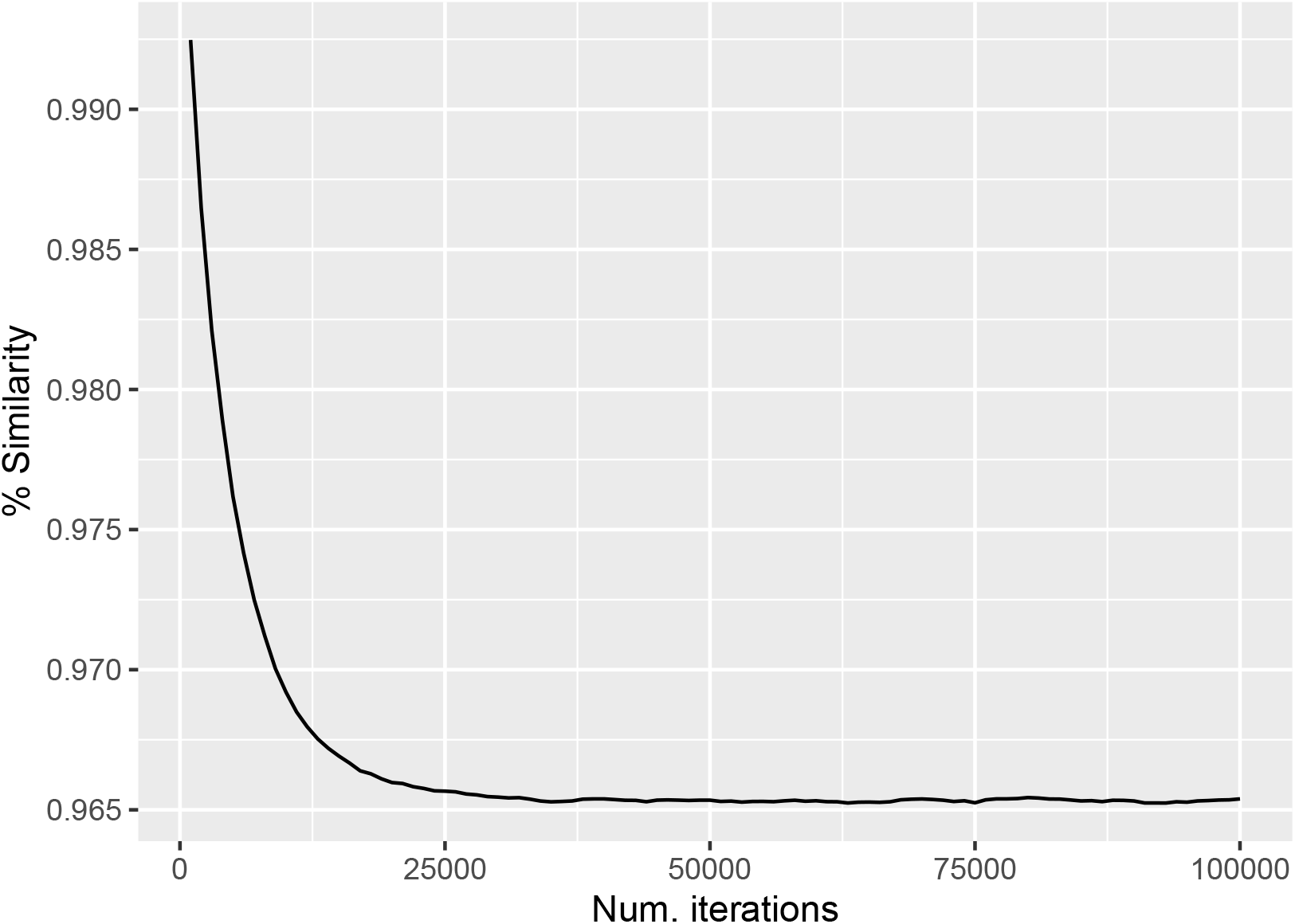
Testing the number of iterations needed to randomize the Australian *Acacia* dataset. The *Acacia* dataset (Mishler et al., 2014) was randomized using the `curveball` algorithm for 100,000 iterations and percentage similarity between the original matrix and the randomized matrix was calculated every 1,000 iterations using the `cpr_iter_sim()` function.

#### cpr_rand_test()

The `cpr_rand_test()` function carries out calculation of observed values, generation of random communities, and calculation of summary statistics as described above in “Analysis workflow”. The main arguments to this function are the input community and phylogeny, type of null model, and settings for the null model. For a full list of null models to choose from, run `?vegan::commsim()`. It should be noted that the type of null model, number of random communities, and number of swapping iterations performed per random community (for swapping algorithms) all may strongly affect results of `cpr_rand_test()` (or any metric that is based on comparison to a set of random communities; Gotelli, 2001). While it is beyond the scope of this paper to provide a full discussion of null models in ecology, we have provided details about how to explore appropriate null model settings with `canaper` in the “How many randomizations?” vignette (https://docs.ropensci.org/canaper/articles/how-many-rand.html).

The output is a data frame with communities as rows and summary statistics in columns. A large number of summary statistics, including all of those needed to calculate CANAPE, are produced. For a full explanation of all output columns, see Appendix S1 or run `?cpr_rand_test()`.

The following code demonstrates usage of `cpr_rand_test()`, using the same example dataset as above.

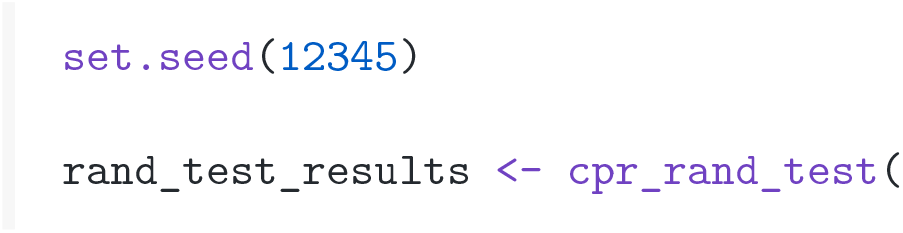
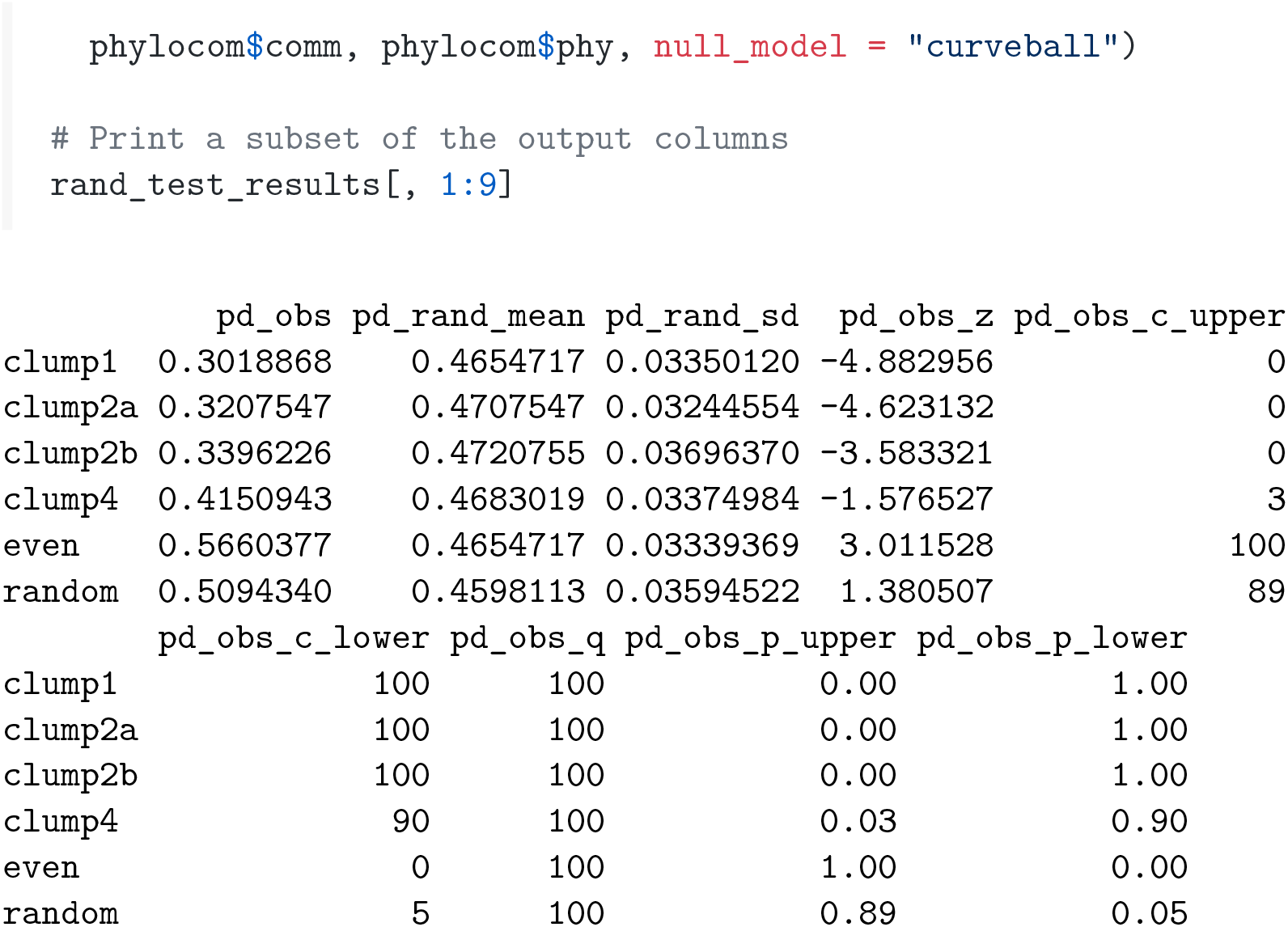

#### cpr_classify_endem()

The `cpr_classify_endem()` function classifies endemism types for the output of `cpr_rand_test()` as described above in “Analysis workflow”. The input is a data frame including the following columns calculated by `cpr_rand_test(): pe_obs_p_upper` (upper *p*-value comparing observed PE to random values), `pe_alt_obs_p_upper` (upper *p*-value comparing observed PE on alternate tree to random values), `rpe_obs_p_upper` (upper *p*-value for RPE), and `rpe_obs_p_lower` (lower *p*-value for RPE). The output is the same data frame, with the column endem_type appended. Values of `endem_type` include paleo (paleoendemic), `neo` (neoendemic), `not significant, mixed` (mixed endemism), and `super` (super-endemic).

The following code demonstrates usage of `cpr_classify_endem()` with the output from `cpr_rand_test()` (note that for this small example, not all possible types of endem_type are produced).

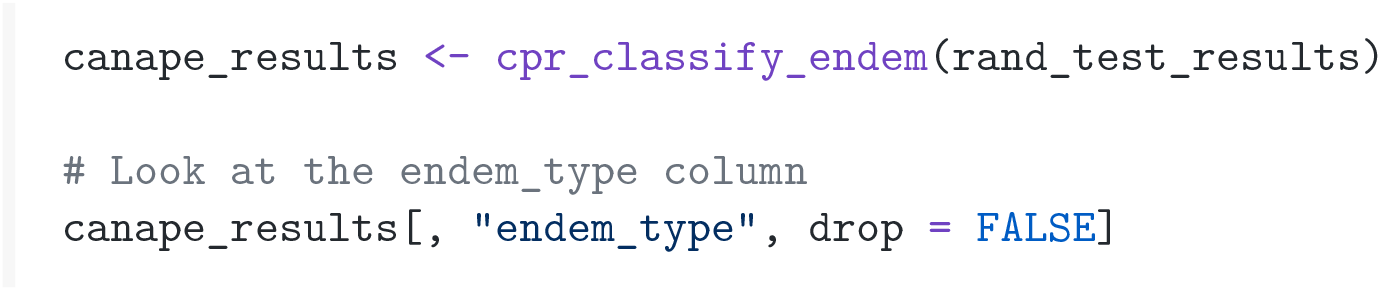
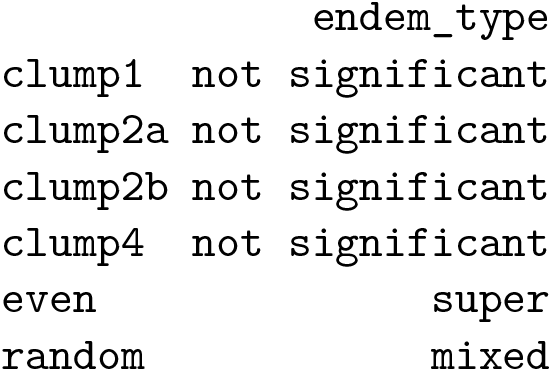

In addition to `cpr_classify_endem(), canaper` includes the `cpr_classify_signif()` function for assessing significance level of PD, RPD, PE, and RPE. Both `cpr_classify_endem()` and `cpr_classify_signif()` take a data frame as their first argument and return a data frame as output, so they are “pipe friendly”, i.e., can be chained together using pipe operators (%>% or |>).

### Parallel computing

Parallel computing is enabled with the `future` package, which has been designed to allow maximum flexibility in the parallel backend selected by the user (e.g., multiple cores on one machine, multiple remote machines, etc.). Parallelization is applied to the calculation of summary statistics for each random community, as there are potentially many random communities (typically > 100 for a robust analysis, though this depends on the dataset). To use parallel computing, no changes are needed for `cpr_rand_test()`etc. Rather, `future` is loaded with `library()`, then a parallel back-end is specified with plan(). The user is advised to consult the `future` website (https://future.futureverse.org/) for more information on specifying a parallel backend.

The following code demonstrates parallel computing.

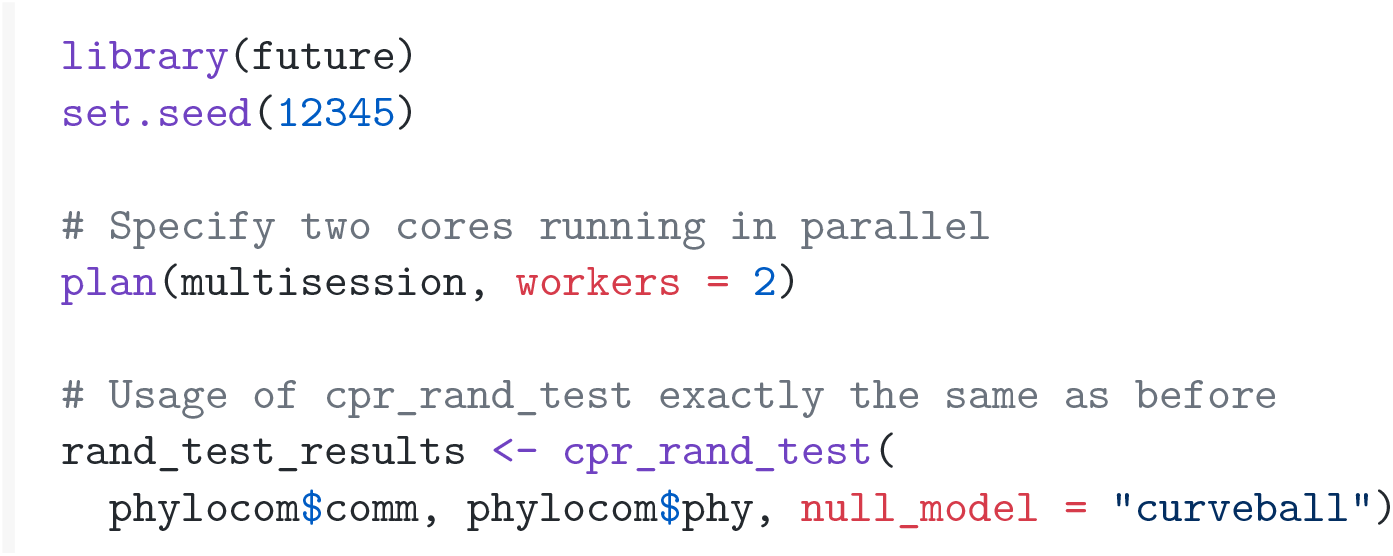

There will not be a noticeable decrease in computing time since this dataset is so small. However, parallelization can greatly decrease computing time for large datasets; more details are available in the “Parallel computing” vignette (https://docs.ropensci.org/canaper/articles/parallel.html).

### Datasets

`canaper` comes with three datasets for testing and demonstration (Table 1). Each is a list with two named elements, `comm` (community matrix, data frame) and `phy` (phylogenetic tree, list of class `“phylo”). acacia` includes data of Australian *Acacia* analyzed by Mishler et al. (2014). `phylocom` and `biod_example` are fictitious datasets compiled for testing software, and were originally published in Phylocom (Webb et al., 2008) and Biodiverse (Laffan et al., 2010), respectively.

### Example: Australian *Acacia*

#### Analysis with canaper

To demonstrate usage with a real dataset, we reproduced the analysis of Mishler et al. (2014), who conducted CANAPE on Australian *Acacia* using Biodiverse. All `canaper` analyses were run with `canaper` v1.0.0 in R v4.2.1. We used the curveball randomization algorithm with 50,000 iterations, since `cpr_iter_sim()` indicated this was sufficient to randomize the matrix (see “Major Functions”). Once the randomization algorithm and its settings have been selected, CANAPE can be conducted with just two commands.

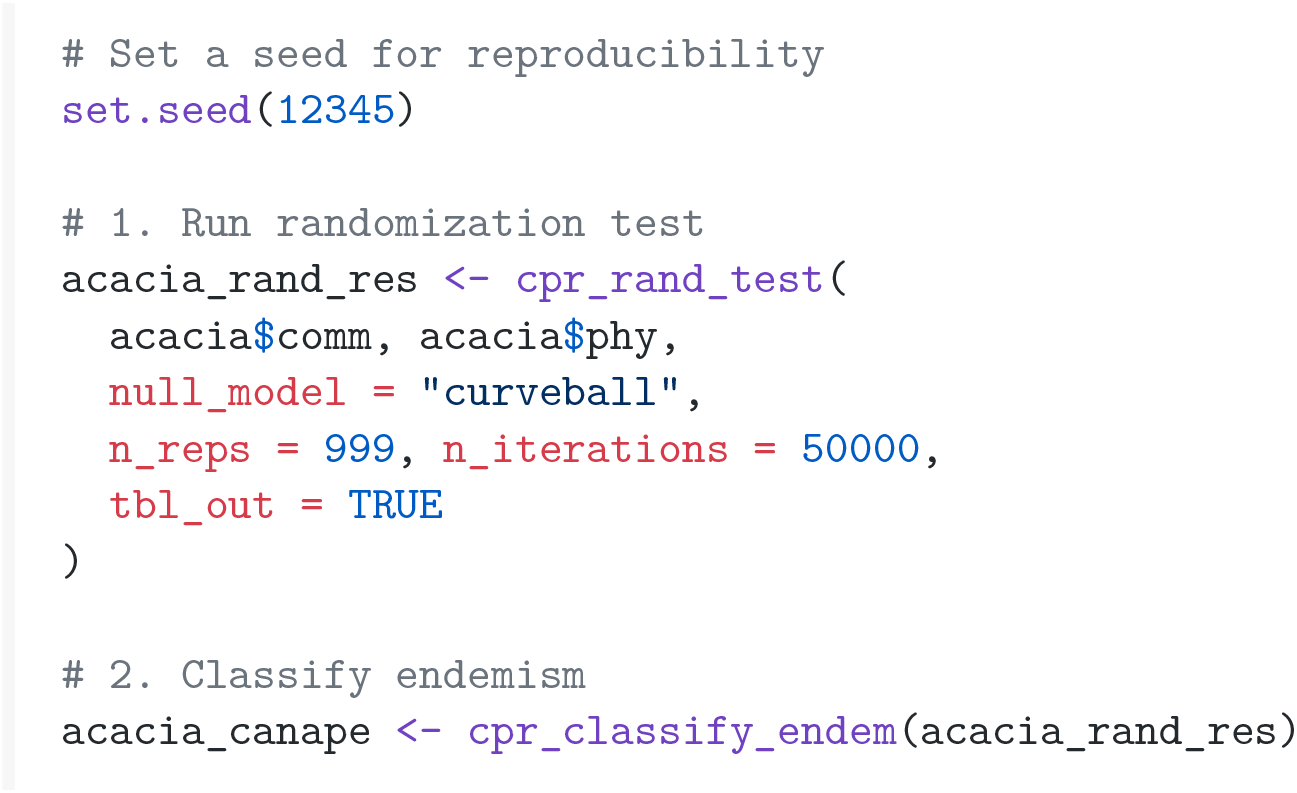

`canaper` does not include any plotting functions to visualize the results. Rather, we recommend the `ggplot2` package or base R graphics to visualize results. Here, we demonstrate use of the `ggplot2` and `patchwork` packages to visualize the output of `canaper` (Fig. 3).

**Fig. 3.**
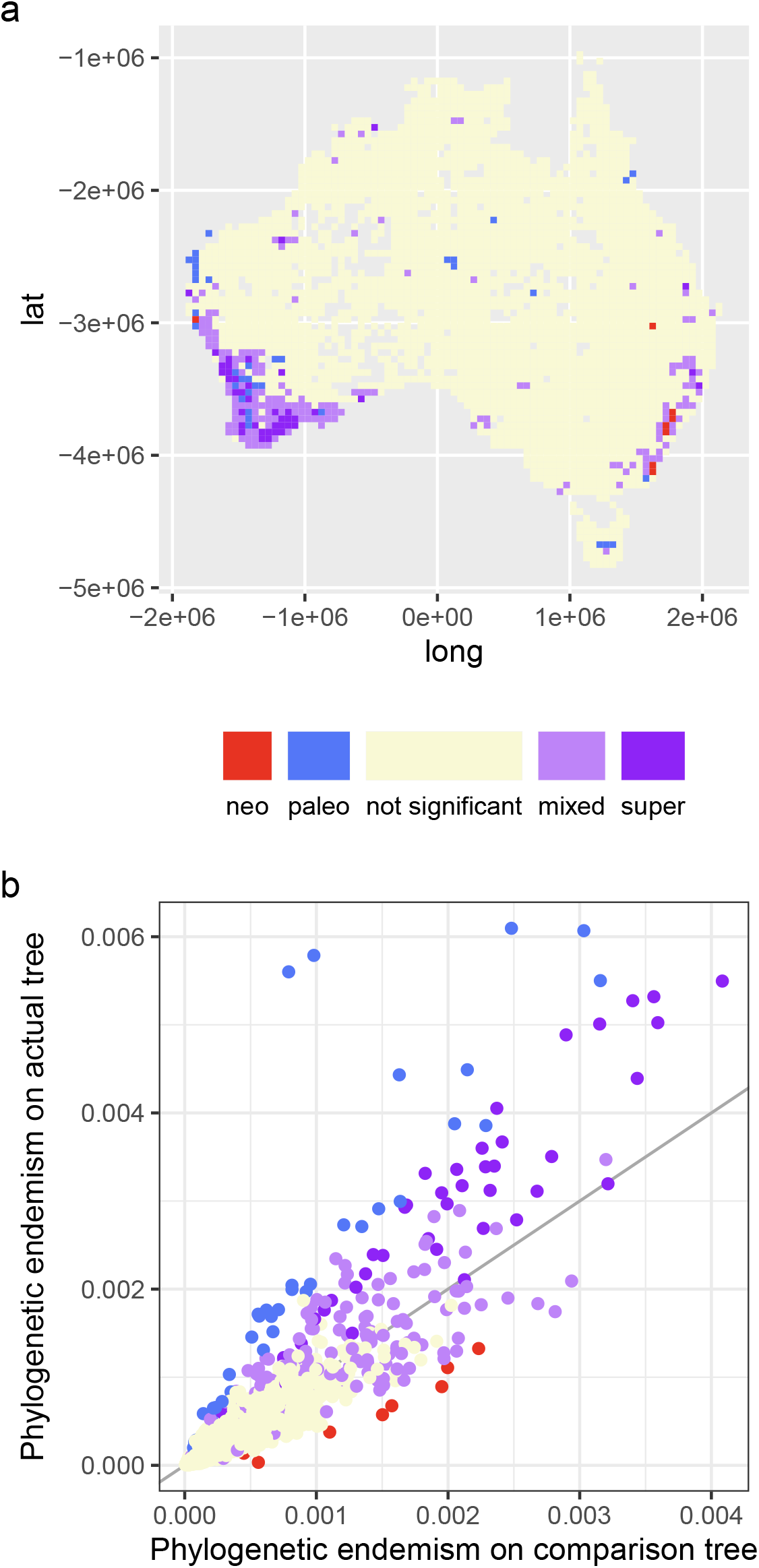
Categorical analysis of neo- and paleo-endemism (CANAPE) of Australian *Acacia*. (a) Map of Australia showing grid-cells (communities) colored by endemism type. Latitude and longitude projected into equal area Australian Albers (EPSG:3577) coordinate system (Butler et al., 2007); units in m. (b) Scatterplot of comparing phylogenetic endemism (PE) of each community as measured on the original tree vs. a comparison tree with all non-zero branch lengths set to equal length, colored according to endemism type. This figure reproduces Figure 3 of Mishler et al. (2014).

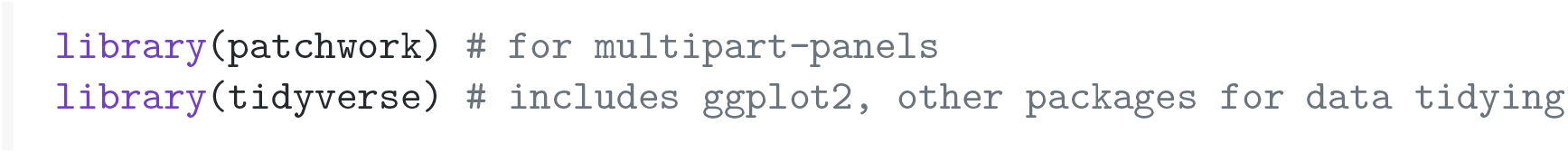
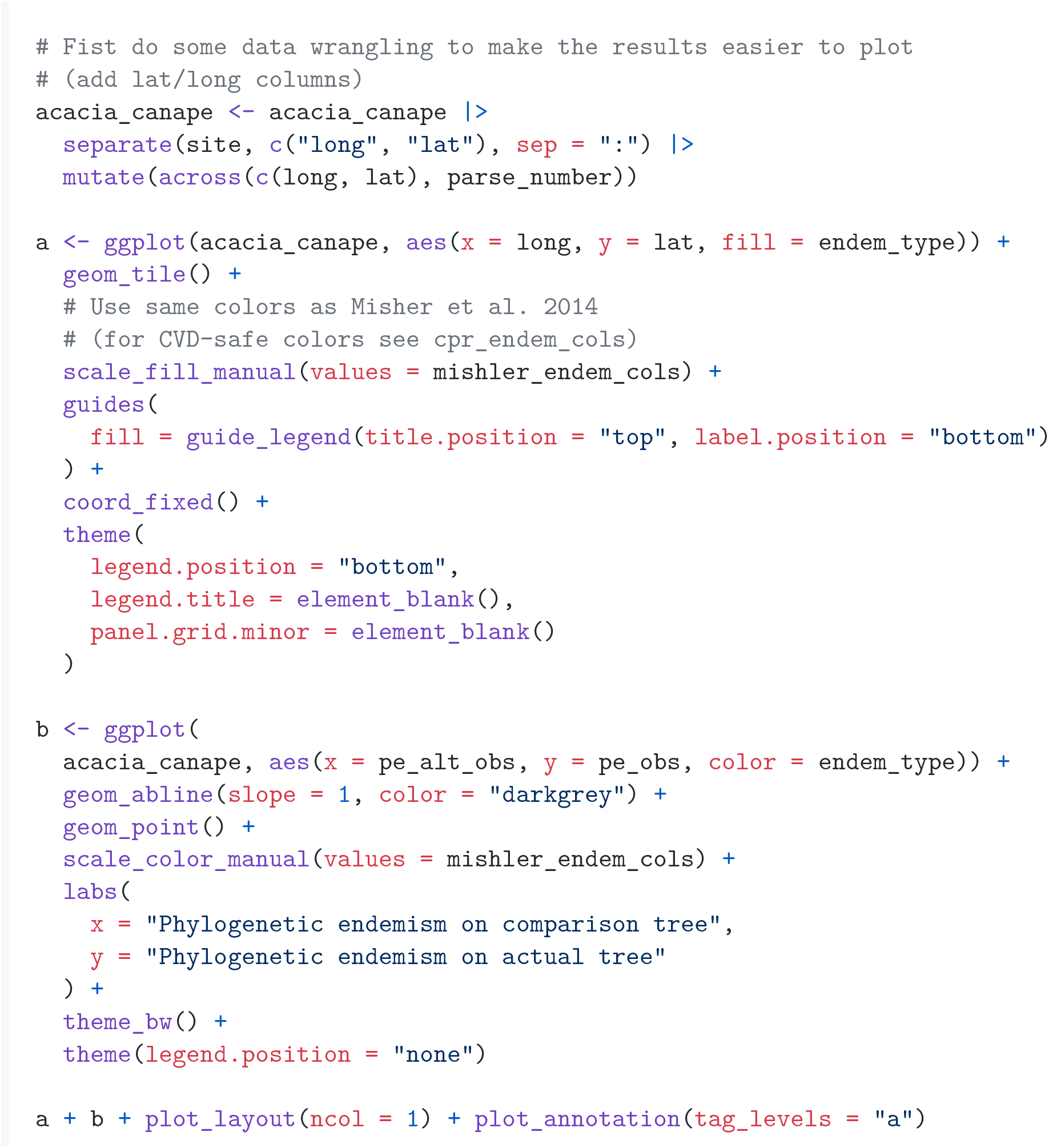

As in Figure 3 of Mishler et al. (2014), grid-cells with significant endemism are primarily located on the coasts, with mostly non-significant grid-cells in the interior (Fig. 3). Furthermore, the endemism types largely correspond between the two figures.

The color palette used in Fig. 3, `mishler_endem_cols`, is the same as that used in Mishler et al. (2014). However, this palette may not be distinguishable to people with color-vision deficiency (CVD). Alternate color palettes (e.g., `cpr_endem_cols`) are also available in `canaper` thatinclude CVD-safe colors (Okabe & Ito, 2002). Palettes for plotting CANAPE and *p*-rank results can be selected with the `cpr_make_pal()` function. The same plot visualized with CVD-safe colors is available in the online supporting material (Fig. S1).

### Comparison with Biodiverse

We also re-ran the *Acacia* analysis using Biodiverse v3.1 with the same settings as Mishler et al. (2014) (999 replicates of the `rand_structured` null model) and compared the results with those from `canaper`. Importantly, there is no expectation that results between the two should match exactly, for two reasons. First, the null model used between the Biodiverse and `canaper` analyses differ (`rand_structured` and `curveball`, respectively). `rand_structured` is not currently available in R, but we hope to add this to a future version of `canaper`. Second, the random communities generated in each run will be different, so the exact *p*-values will also be different. With a sufficiently high number of random communities, significance (e.g., at the *α* = 0.5 level) is expected to converge, but there may be borderline cases that appear significant in some analyses and non-significant in others.

When we compared endemism type between the `canaper` and Biodiverse CANAPE results for Australian *Acacia*, they agreed in 98.4% of grid-cells (Table 2). The total number of significant cells was very similar (*n* = 257 and 253 sites, `canaper` and Biodiverse, respectively). The `rand_structured` and `curveball` null models both preserve total abundance per species and richness of each site (community matrix marginal sums); in that sense, they are relatively conservative null models (Strona et al., 2018), which may explain the relative high agreement between results. Selection of an appropriate null model is beyond the scope of this paper, but must be considered carefully in any community ecological analysis.

**Table 2:** Comparison of CANAPE results for Australian *Acacia* between `canaper` and Biodiverse

| Endemism type | $n$ sites (`canaper`) | $n$ sites (Biodiverse) |
| --- | --- | --- |
| mixed | 172 | 170 |
| neo | 8 | 7 |
| not significant | 2780 | 2784 |
| paleo | 36 | 35 |
| super | 41 | 41 |

Computations were carried out on an MacBook Pro (2019) with 16 GB RAM and a 1.7 GHz, four core processor. Approximate compute times were 44.4 min for Biodiverse and 8.6 min for `canaper` (sequential, i.e., non-parallel mode, for both). However, when parallel computing was enabled for `canaper` with two cores, compute time dropped to 4.9 min. This demonstrates that `canaper` can efficiently compute moderately sized datasets with a personal laptop computer.

### Comparison with other R packages

We are not aware of any other R packages that conduct the entire CANAPE pipeline automatically. However, there is a large number of packages for analyzing species diversity (approximately 40 packages out of 15,300 as of 2019; Pavoine, 2020). Some of those that are more closely related to `canaper` include the following. `picante` (Kembel et al., 2010) was one of the first packages to offer calculation of phylogenetic alpha diversity such as MPD, MNTD, and PD (Webb, 2000). `phyloregion` (Daru et al., 2020) implements sparse matrix encoding to increase computing efficiency of PD, and is used by `canaper. vegan` (Oksanen et al., 2017) performs a wide range of non-phylogenetic community ecology analyses. `Vegan` includes by far the greatest number of algorithms for generating random communities and is used by `canaper. adiv` (Pavoine, 2020) provides a flexible framework for analyzing alpha and beta diversity of biological communities based on either phylogenetic or trait distances. `PhyloMeasures` (Tsirogiannis & Sandel, 2016) implements efficient routines for calculating phylogenetic diversity metrics. `EcoSimR` (Gotelli & Ellison, 2013) provides randomization algorithms as well as functions for characterizing community matrices and null distributions. As of writing, `PhyloMeasures` and `EcoSimR` had been removed from CRAN and may not be under active development.

## Conclusions

The `canaper` package enables CANAPE completely within R for the first time. This will simplify workflows and facilitate reproducibility for researchers studying spatial biodiversity with R. Furthermore, `canaper` features efficient computing routines and simple yet flexible implementation of parallel computing, thereby decreasing computation time. `canaper` has already been used at least three studies while in development (Ellepola et al., 2022; Lu et al., 2022; Nitta et al., 2022). We expect `canaper` will become a major tool in the toolkit of the emerging field of spatial phylogenetics alongside Biodiverse.

## Acknowledgements

This research supported in part by Japan Society for the Promotion of Science (Kakenhi) Grant numbers 16H06279, 22H04925, and 22K15171. Members of the Iwasaki Lab (The University of Tokyo) provided helpful comments while the package was in development. Luis Osorio and Klaus Schliep reviewed the code as part of submission to rOpenSci.

## Author contributions

Joel H. Nitta conceived the ideas and designed methodology; Joel H. Nitta and Shawn W. Laffan analyzed the data; Joel H. Nitta led the writing of the manuscript; Brent D. Mishler edited the manuscript; Wataru Iwasaki provided resources and supervised the study. All authors contributed critically to the drafts and gave final approval for publication.

## Data availability

The `canaper R` package is distributed by the Comprehensive R Archive Network (CRAN), with source code available on GitHub (https://github.com/ropensci/canaper) and Zenodo (https://doi.org/10.5281/zenodo.5094032). Usage of `canaper` is documented at https://docs.ropensci.org/canaper/. Code and data used to conduct analyses and generate this manuscript are available at https://github.com/joelnitta/canaper_ms, and can be run using the Docker image `joelnitta/canaper_ms` available at https://hub.docker.com/r/joelnitta/canaper_ms. The acacia dataset was originally published by Mishler et al. (2014).

## Boxes

Box 1: Glossary of terms.

- Community: A co-occurring set of species.
- Grid-cell: One of a set of spatial units each with equal area that together cover a region of interest; each grid-cell comprises a community. Often used synonymously with “site”.
- Community matrix: A 2-dimensional dataset with communties (grid-cells) on one axis and species on the other. `canaper` expects communities as rows and species as columns.
- Phylogenetic diversity (PD): The total branch length (including the root) connecting the species occurring in a given area (community or grid-cell), measured on an overall tree that includes all the species in the study (Faith, 1992).
- Phylogenetic endemism (PE): Range weighted PD where the weight for each branch is the fraction of its overall range represented in the sample area (typically a grid-cell) (Rosauer et al., 2009).
- Relative phylogenetic diversity (RPD): The ratio of PD measured on the original tree vs. PD measured on a comparison tree with all branch lengths transformed to equal length (Mishler et al., 2014).
- Relative phylogenetic endemism (RPE): The ratio of PE measured on the original tree vs. PE measured on a comparison tree with all branch lengths transformed to equal length (Mishler et al., 2014).
- Neoendemic: An area with a high concentration of short, range-restricted branches; may be due to processes such as recent speciation (radiation).
- Paleoendemic: An area with a high concentration of long, range-restricted branches; may be due to processes such as extinction or colonization by distantly relatives from outside the study area.
- Mixed endemic: An area with a mixture of short and long, range-restricted branches; may be due to multiple processes.
- Super endemic: An area with mixed endemism that is statistically highly significant.

