## Supplemental Information for "canaper: Categorical analysis of neo- and paleo-endemism in R"

**Supporting information**

Joel H. Nitta Shawn W. Laffan Brent D. Mishler
Wataru Iwasaki

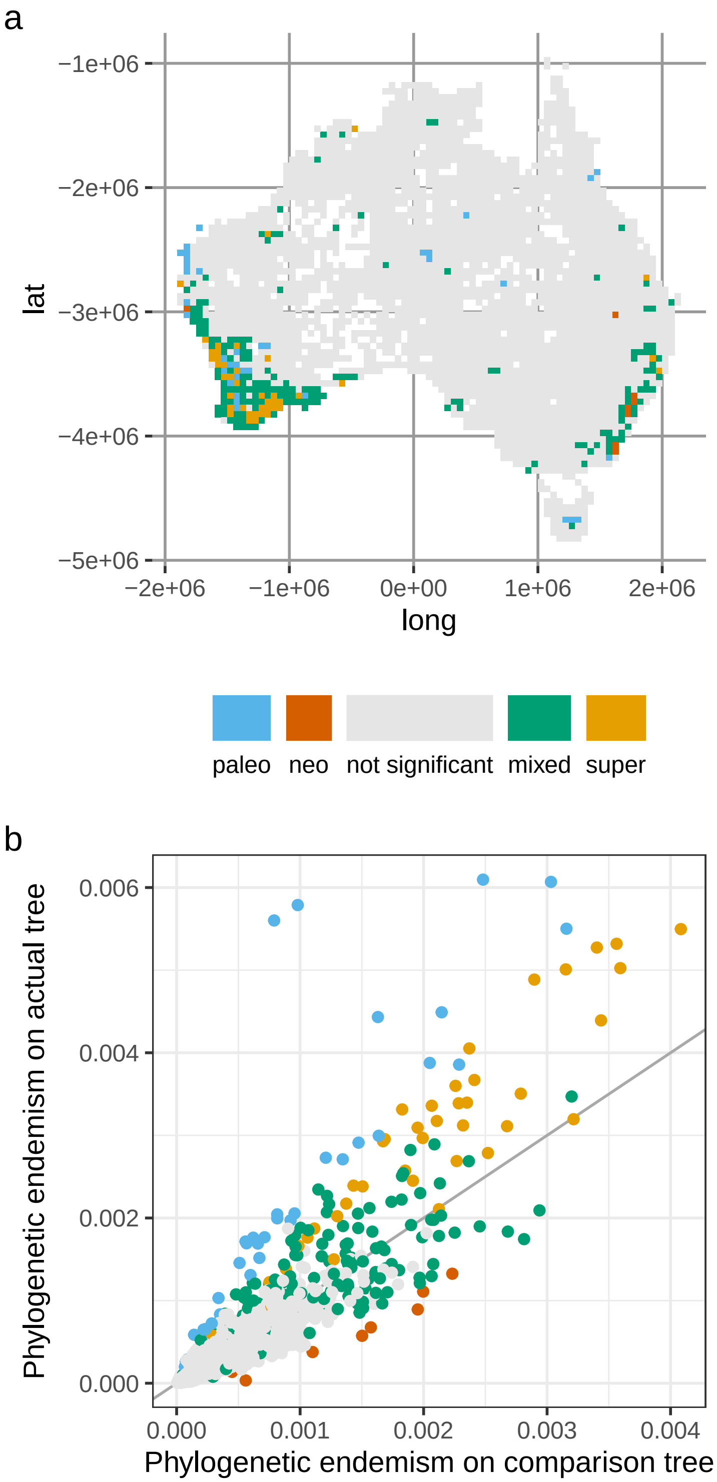


**Fig. S1**: Categorical analysis of neo- and paleo-endemism (CANAPE) of Australian *Acacia*, plotted with color vision deficiency (CVD)-safe colors. (a) Map of Australia showing grid-cells (communities) colored by endemism type. Latitude and longitude projected into equal area Australian Albers (EPSG:3577) coordinate system; units in m. (b) Scatterplot of comparing phylogenetic endemism (PE) of each community as measured on the original tree vs. a comparison tree with all non-zero branch lengths set to equal length, colored according to endemism type.

**Appendix S1**: Columns in data frame produced by `cpr_rand_test()`.

`cpr_rand_test()` calculates six basic metrics for each grid-cell:

- `pd`: phylogenetic diversity (PD).
- `pd_alt`: alternative PD (PD measured on comparison tree with all branch lengths transformed to equal length).
- `pe`: phylogenetic endemism (PE).
- `pe_alt`: alternative PE (PE measured on comparison tree).
- `rpd`: relative PD (ratio of `pd` to `pd_alt`).
- `rpe`: relative PE (ratio of `pe` to `pe_alt`).

For each of the six basic metrics, nine columns (54 total) are output by `cpr_rand_test()`:

- `*_obs`: Observed value.
- `*_obs_c_lower`: Count of times observed value was lower than random values.
- `*_obs_c_upper`: Count of times observed value was higher than random values.
- `*_obs_p_lower`: Percentage of times observed value was lower than random values.
- `*_obs_p_upper`: Percentage of times observed value was higher than random values.
- `*_obs_q`: Count of the non-NA random values used for comparison.
- `*_obs_z`: Standard effect size ($z$-score).
- `*_rand_mean`: Mean of the random values.
- `*_rand_sd`: Standard deviation of the random values.

So for `pd` the output columns would include `pd_obs`, `pd_obs_c_lower`, etc. The $p$-values (`*_obs_p_upper`, etc.) are then used to categorize endemism type according to CANAPE (Appendix S2).

To minimize memory requirements, the individual random communities are not stored in memory when calculating metrics. Since hundreds to thousands or more random replicates may be required, the memory usage could be quite high for large matrices if each replicate was stored. Rather, as each random community is generated, the summary statistics are computed immediately and stored (the same approach is used in Biodiverse).

**Appendix S2**: Criteria for categorical analysis of neo- and paleo-endemism (CANAPE).

Significance assessed at $\alpha = 0.05$ (significant) or $\alpha = 0.01$ (highly significant). Metrics described in Appendix S1. Not all studies include super-endemism.

- If either `pe_obs_p_upper` or `pe_alt_obs_p_upper` are significantly high (one-sided test), then look for paleo- or neo-endemism.
  - If `rpe_obs_p_upper` is significantly high (two-sided test), then grid-cell is paleo-endemic.
  - Else if `rpe_obs_p_lower` is significantly low (two-sided test), then grid-cell is neo-endemic.
  - Else there is mixed age endemism, in which case:
    - If both `pe_obs_p_upper` and `pe_alt_obs_p_upper` are highly significant ($p < 0.01$), then grid-cell is super-endemic.
    - Else grid-cell is mixed-endemic.
- Else if neither `pe_obs_p_upper` nor `pe_alt_obs_p_upper` are significantly high, grid-cell is non-endemic.
